# OpenTn5: Open-Source Resource for Robust and Scalable Tn5 Transposase Purification and Characterization

**DOI:** 10.1101/2024.07.11.602973

**Authors:** Jan Soroczynski, Lauren J. Anderson, Joanna L. Yeung, Justin M. Rendleman, Deena A. Oren, Hide A. Konishi, Viviana I. Risca

## Abstract

Tagmentation combines DNA fragmentation and sequencing adapter addition by leveraging the transposition activity of the bacterial cut-and-paste Tn5 transposase, to enable efficient sequencing library preparation. Here we present an open-source protocol for the generation of multi-purpose hyperactive Tn5 transposase, including its benchmarking in CUT&Tag, bulk and single-cell ATAC-seq. The OpenTn5 protocol yields multi-milligram quantities of pG-Tn5^E54K, L372P^ protein per liter of *E. coli* culture, sufficient for thousands of tagmentation reactions and the enzyme retains activity in storage for more than a year.

## Introduction

Tn5 is a bacterial cut-and-paste transposase, which can simultaneously fragment target DNA and append its ends with custom DNA adapter sequences in a process referred to as tagmentation[1] (Fig. 1A). In the reaction, Tn5 homodimer first binds two double-stranded DNA molecules encoding the mosaic end sequence[2], referred to as MEDS adapters[3], to form a transposome competent for target DNA binding. In the presence of Mg^2+^, the transposome-target synapse complex then performs a strand transfer reaction, in which the 3’OH groups of the MEDS adapters perform a nucleophilic attack on the target DNA backbone offset by 9 base pairs. The resulting target DNA ends are now appended with adapter sequences and contain 9 bprecessed 3’OH ends. This simultaneous target DNA cleavage and adapter attachment activity of Tn5 is referred to as tagmentation. Through decades of research pioneered by William S. Reznikoff and colleagues, many gain-of-function Tn5 mutations have been isolated that greatly increase the rate of Tn5 transposition, and are commonly referred to as hyperactive mutants[1]. Two of these, E54K[4] and L372P[5], are sufficient for a dramatic increase in activity and are the basal hyperactive mutations historically used in tagmentation applications. Subsequent studies have also investigated the effects of additional Tn5 mutations on further augmentation of tagmentation activity[6], as well as altering target DNA sequence insertion bias[7–9].

**Figure 1:**
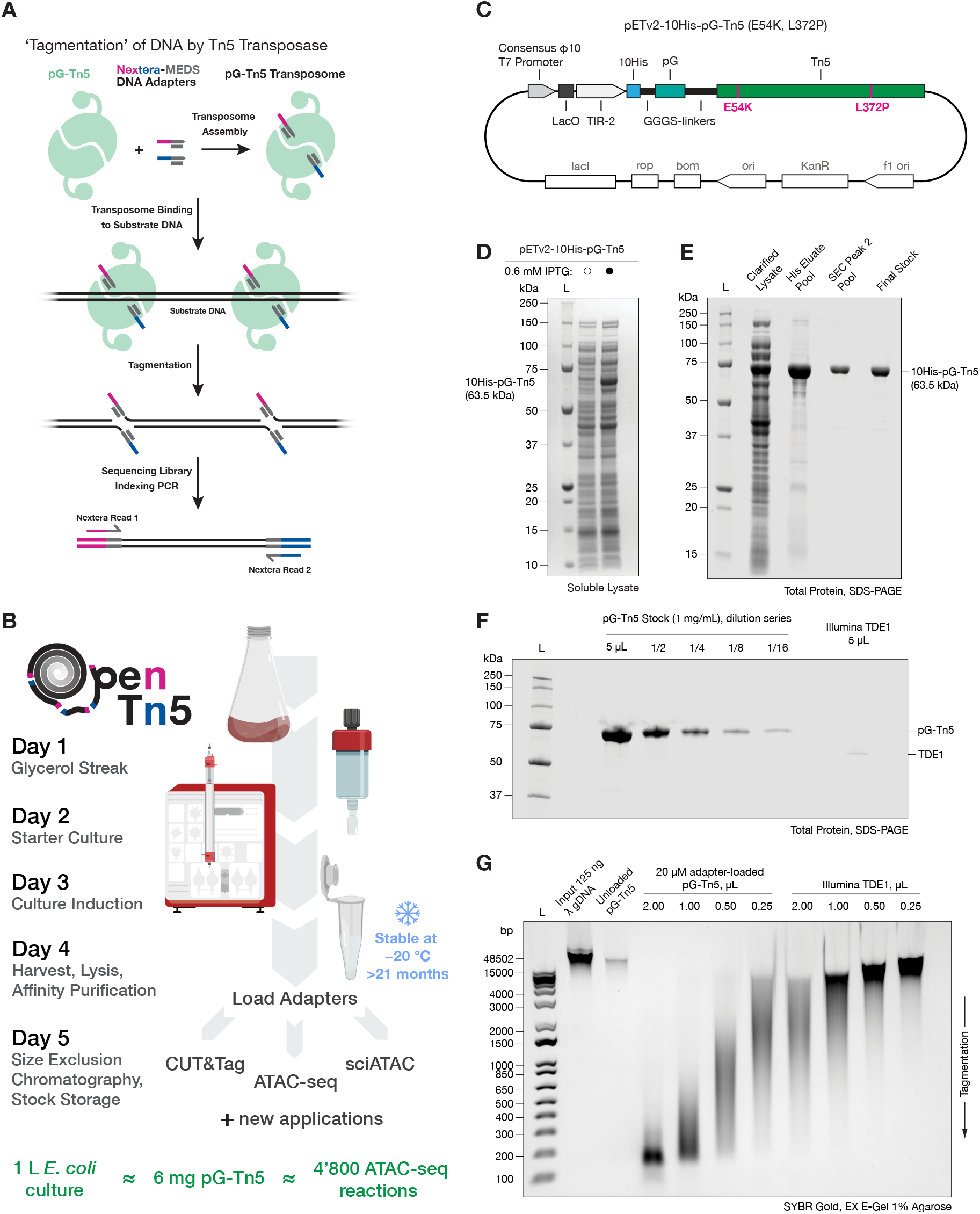
OpenTn5 enables robust, scalable, open-source Tn5 transposase purification for multiple applications. **A**. Diagram of simultaneous DNA fragmentation and addition of DNA sequencing adapters *in vitro* by Tn5 transposition reaction, referred to as tagmentation. **B**. Expression and purification of pG-Tn5 can be carried out within 5 days and yields multi-milligram amounts of enzyme, sufficient for thousands of tagmentation reactions. Less than 1.25 µg of pG-Tn5 is required per 50’000 cell input ATAC-seq reaction. **C**. Schematic of 10His-pG-Tn5(E54K, L372P) expression plasmid, VR124 (Addgene #198468). **D**. Robust over-expression of soluble 10His-pG-Tn5(E54K, L372P) protein. **E**. His-tag affinity and size exclusion chromatography (SEC) enable efficient purification of pG-Tn5 homodimer. **F**. Comparison of pG-Tn5 adapter unloaded stock at 1 mg/mL with commercial Illumina TDE1 enzyme (Cat #20034198). **G**. Comparison of tagmentation activity of pG-Tn5 (1 mg/mL) and Illumina TDE1 using equal transposome volumes on Lambda genomic DNA (λ gDNA).

Tagmentation has been leveraged for streamlined preparation of high-throughput DNA sequencing libraries from sample DNA using custom sequencing adapter sequences. The efficiency and versatility of Tn5 tagmentation has given rise to a variety of genomics technologies for investigating chromatin organization, transcription and cell identity, with innovations continuing to be made in novel Tn5 protein fusions as well as custom MEDS adapter designs[10–19]. Despite the accomplishments of tagmentation-based methods to date, in-house production of high-quality Tn5 enzyme remains a persistent challenge[10] which hinders both ongoing efforts in the development of novel tagmentation applications and the equitable, broad adoption of existing methods.

Here we present our current integration of optimizations into a “start-to-finish”, open-source protocol for the generation of a robust, all-purpose hyperactive Tn5 transposase, including validation across multiple sequencing-based methods that rely on transposition[9, 20, 21]. We demonstrate the functional equivalence of the produced pG-Tn5^E54K, L372P^ to commercially-sourced Tn5 used for CUT&Tag, but importantly we show the same enzyme can also be readily used for assays not requiring protein G for antibody-directed recruitment of Tn5, such as bulk and single-cell ATAC-seq, underscoring its versatility. Notably, pG-Tn5^E54K, L372P^ can be readily loaded with custom DNA adapters. We hope that this OpenTn5 protocol will further disseminate existing applications of tagmentation, as well as facilitate future method development efforts.

## Results

We sought to develop a robust protocol for production of high quality, hyperactive Tn5 which could be reproducibly carried out using generally available molecular biology laboratory equipment. To this end, the OpenTn5 protocol can be carried out within one work week (Fig. 1B). At 1x scale, the protocol uses 500 mL *E. coli* culture to yield roughly 3 milligrams of final pure pG-Tn5^E54K, L372P^ homodimer stock (Fig. S1A-F), which is stable for > 21 months at −20 °C (Fig. S1G-H). This amount is sufficient for several thousand reactions, e.g. one 50’000 cell input ATAC-seq reaction consumes 1.25 µg of pG-Tn5^E54K, L372P^.The transposase stock is stored unloaded, and adapter loading is carried out at time of use, ensuring stock stability and allowing versatility for different tagmentation applications.

We provide a detailed step-by-step protocol, including an assay for measuring Tn5 tagmentation activity as supplementary materials. The expression plasmid is available from Addgene in two *E. coli* strains DH5α (#198467) and NEB T7 Express *lysY/I*^*q*^ (#198468), for plasmid propagation and protein expression, respectively.

### *E. coli* expression, purification and storage of 10HispG-Tn5^E54K, L372P^

We began by focusing on optimizing the expression and purification of the 10His-pG-Tn5^E54K, L372P^ construct previously described by Xu and colleagues[21], which showed good over-expression in *E. coli*. Xu and colleagues demonstrated that Tn5 does not tolerate Cterminal tagging[21], likely due to the sensitivity of the nearby dimerization domain that is critical for transposition[22]. We note that the pTXB1-derived Tn5 expression plasmids[3] employ a C-terminal intein tag, which may contribute to the difficulty in obtaining active Tn5 protein using these constructs. The N-terminal 10His-protein G tag we use serves to promote solubility, facilitates straightforward affinity purification, and can be used for antibody-mediated Tn5 recruitment in CUT&Tag. We cloned an *E. coli* codon-optimized 10HispG-Tn5^E54K, L372P^ protein coding sequence into an optimized derivative of IPTG-inducible pET28a protein expression vector described by Shilling and colleagues[20] (Fig. 1C). We refer to this expression plasmid as pETv2-10His-pG-Tn5 ^E54K, L372P^.

The resulting construct showed robust IPTG-dependent over-expression of soluble 10His-pG-Tn5 at 18 °C when introduced into the NEB T7 Express *lysY/I*^*q*^ *E. coli* strain. Notably, we did not encounter problems with plasmid toxicity in *E. coli* neither during cloning in the NEB 5-alpha strain nor during protein expression. Crucially, the majority of the over-expressed protein reliably remained in the soluble lysate fraction (Fig. 1D). We optimized *E. coli* culture conditions; using richer Terrific Broth (TB) instead of Lysogeny Broth (LB) media[23], supplemented with an anti-foam agent in double-baffled 2 L flasks to maximize the culture aeration. These changes further improve the soluble expression yield independently of construct design, and maximize the *E. coli* biomass harvested, reducing culture volumes required to be handled.

We use a standard sonication lysis without adding exogenous nucleases to minimize the risk of their carryover throughout and adapted a standard immobilized metal affinity chromatography (IMAC), specifically Ni-NTA resin affinity purification without subsequent cleavage of the 10His-pG tag, which streamlines the protocol. We extensively wash the resin-bound Tn5 using a high salt buffer to minimize non-specific binding of *E. coli* nucleic acid contaminants to Tn5 (Fig. S1A). We do not routinely perform any additional nucleic acid removal during the purification, such as polyethylenimine (PEI) precipitation, ion or heparin exchange, as in our experience the trace amounts of *E. coli* genomic DNA impurities in the final Tn5 stock do not significantly interfere in typical sequencing assay applications (Fig. S2B).

We polish the Ni-NTA column eluate using size exclusion chromatography (SEC), following the protocol developed by Hennig and colleagues[9] (Fig. 1E). We confirmed that SEC successfully separates full-length 10His-pG-Tn5 homodimers away from Tn5 aggregates, heterodimers containing truncated Tn5 and spuriously cleaved 10His-pG tag (Fig. S1B). We pool the SEC fractions corresponding to the 10His-pG-Tn5 homodimer peak and concentrate them to a concentration of 2 mg/mL in the SEC buffer. We prepare the final Tn5 stock by adding an equal volume of pure glycerol to the concentrated protein (1 mg/mL protein in 55% glycerol, final) and store the liquid stock at −20 °C in enzyme cooler blocks. We see a negligible loss of 10His-pG-Tn5 activity after almost 2 years of storage under these conditions (21 months, (Fig. S1G)). We do not load Tn5 with MEDS DNA adapters prior to storage, as we have found the assembled transposomes to be unstable. This is likely due to the change in the effective final buffer composition resulting from diluting the protein with adapters resuspended in a low-salt buffer.

We believe the SEC step, often omitted from many existing Tn5 purification protocols, is likely critical to the long-term stability of Tn5 as the aggregates present in the final protein glycerol stock may seed further aggregation and thereby lead to enzyme deterioration[24]. Additionally, SEC purification is likely important for reproducible performance in antibody-based tagmentation assays in which an excess of 10His-pG proteolytic degradation fragments may significantly compete with 10His-pG-Tn5 for antibody binding, thereby reducing assay sensitivity.

### Benchmarking 10His-pG-Tn5 ^E54K, L372P^

#### *in vitro* activity

We compared the protein concentrations of our 1 mg/mL pG-Tn5 stock against commercially available Illumina TDE1 enzyme (CAT: #20034198, LOT: 20541598), provided at an unspecified concentration. We loaded equal volumes (5 µL) of both enzymes, alongside a dilution series of the adapter-unloaded 1 mg/mL pG-Tn5 stock and measured total protein content by SDS-PAGE followed by Coomassie staining (Fig. 1F). Illumina TDE1 protein concentration was roughly 16-fold lower than that of the 1 mg/mL pG-Tn5 stock, giving an estimate of 62.5 µg/mL protein concentration of the adapter pre-loaded commercial TDE1 enzyme.

To measure the tagmentation activity of Tn5, we use commercially available Lambda phage genomic DNA (λ gDNA) as substrate. λ gDNA serves as a reproducible assay substrate; it is supplied as double-stranded, high-molecular weight (48.5 kb) DNA. The large size of λ gDNA allows for a sensitive, yet simple readout of tagmentation activity by gel electrophoresis (Fig. 1G, Fig. S1G). We load the pG-Tn5 stock with an equal volume of 20 µM Nextera MEDS adapters, corresponding to a modest stoichiometric excess relative to Tn5, to ensure saturated adapter loading (see Supplementary Protocol for details on adapter loading and tagmentation assay). The resulting adapter- loaded pG-Tn5 transposome protein concentration is half that of unloaded stock (0.5 mg/mL). As noted above, we load pG-Tn5 with adapters on a single-use basis to maintain long-term activity in storage.

We compared the tagmentation activity of matched-volume serial dilutions of pG-Tn5 and TDE1 transposomes using the Lambda assay, which showed that 0.5 mg/mL pG-Tn5 transposome has an approximately 8fold higher tagmentation activity than Illumina TDE1, on volume-to-volume basis (Fig. 1G). This 8-fold difference is therefore within the magnitude expected based on the 16-fold difference in Tn5 protein stock concentration, as determined by SDS-PAGE (Fig. 1F), followed by a twofold pG-Tn5 dilution upon adapter loading.

#### CUT&Tag

To determine if the performance of in-house pG-Tn5 is comparable to commercial pAG-Tn5 from Epicypher (SKU: 15-1017), we performed CUT&Tag using both enzymes to profile the distribution of H3K27me3 and H3K27ac histone modifications in MCF7 cells. Two replicates per condition were generated to assess reproducibility. After confirming that the peaks called were highly reproducible across replicate and enzyme, Spearman correlation coefficients were calculated on the rlog normalized counts matrix. The sample correlation matrix shows that samples cluster by histone modifications and different enzymes for the same histone modification clustered with each other (Fig. 2A). The correlation coefficient within each histone modification was between 0.75-1.0. Genome wide coverage tracks also show a similar pattern of distribution across different Tn5 enzymes for each histone modification (Fig. 2B).

**Figure 2:**
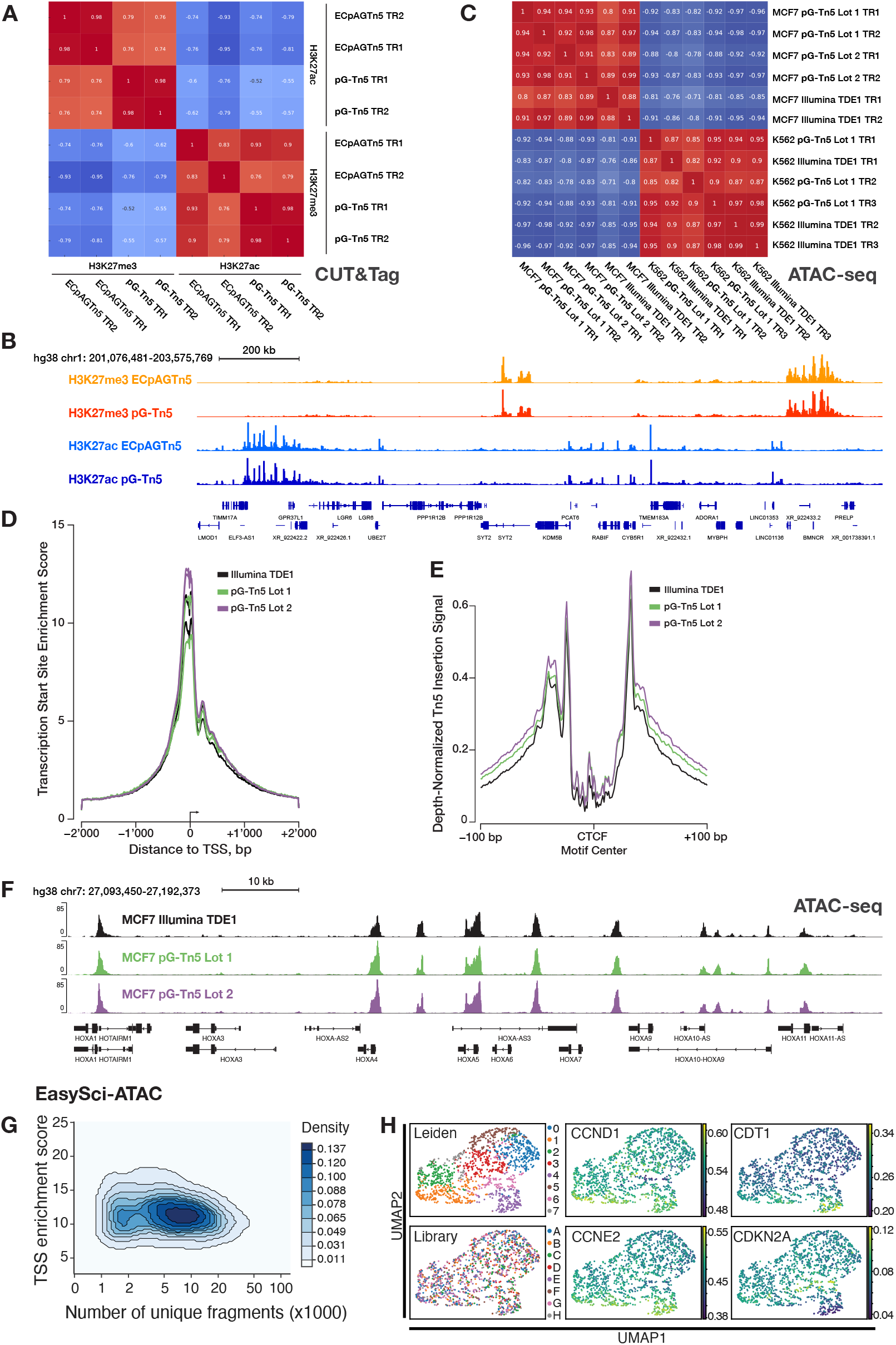
Application and validation of pG-Tn5 in CUT&Tag, ATAC-seq and EasySci-ATAC assays. **A**. Spearman rank correlation of anti-H3K27me3 and anti-H3K27ac CUT&Tag signal under common peaks generated using pG-Tn5 and Epicypher pAG-Tn5 (ECpAGTn5) enzyme (Cat #15-1017) in human MCF7 cells. **B**. Genome track visualization of combined two technical replicates of above conditions, as in panel A. H3K27me3 and H3K27ac signal from pG-Tn5 and pAG-Tn5 experiments plotted on same sequencing-depth scaled y-axes, respectively. **C**. Spearman rank correlation of signal under common peaks, generated using ATAC-seq protocol generated libraries from human K562 and MCF7 cells using pG-Tn5 and Illumina TDE1, using matched transposome volumes. **D**. Transcription start site enrichment score of ATAC-seq signal in human MCF7 cells, comparison of two purification lots of pG-Tn5 and Illumina TDE1 enzyme, two technical replicates each. **E**. Sequencing-depth normalized Tn5 insertion signal around occupied CTCF sites in human MCF7 cells for ATAC-seq signal from pG-Tn5 and Illumina TDE1, as above in panel D. **F**. Genome track visualization of ATAC-seq signal from pG-Tn5 and Illumina TDE1 generated libraries, combined two technical replicates of above conditions, as in panel D and E. **G**. Single-cell accessible chromatin mapping using pG-Tn5 loaded with six unique barcoded adapter combinations, using the EasySci-ATAC protocol in human MCF7 cells. TSS enrichment scores for 2’672 single cells, majority of the cells showing score >10. **H**. Leiden clustering annotations (0-7) of EasySci-ATAC data from cycling human MCF7 cells from eight library replicates (A-H), showing no library-based bias in clustering. Signal values from several cell cycle-associated genes is shown.

#### ATAC-seq

To compare the performance of pG-Tn5 with commercially available Tn5 (Illumina TDE1, CAT: #20034198), we conducted ATAC-seq experiments in MCF7 and K562 human cell lines with matched enzyme volumes. Both enzymes generated high-quality ATAC-seq libraries with similar performance (Supplementary Table 1, Fig. S2A-D). Libraries from the same cell type were highly correlated by peaks and show highly similar genomic coverage (Fig. 2C, F). TSS Enrichment Scores, commonly used as a signal-to-noise QC metric, were comparable across replicates (Fig. 2D). In MCF7 cells, the average TSS scores were 10.9 with Illumina TDE1 and 11.5 with pG-Tn5 (Fig. 2D). In K562 cells, the scores were 12.7 with Illumina TDE1 and 11.7 with pG-Tn5 (Fig. S2C). CTCF site insertion and coverage profiles were highly similar, indicating that protein G does not appreciably sterically interfere with Tn5 transposition, making pG-Tn5 suitable for investigating regulatory landscapes(Fig. 2E, Fig. S2D).

We performed a pG-Tn5 transposome titration in K562 cells to understand how varying pG-Tn5 enzyme concentration influences ATAC-seq signal globally, over 1 kb-sized genomic bins (see Methods). We found samples tagmented with 2.5, 5.0 and 7.5 µL of pG-Tn5 were most correlated, with samples made with 1 µL pG-Tn5 to be most correlated with Illumina TDE1 (Fig. S2A). TSS scores remained high even when using as little as 1 µL of pG-Tn5 S2C).

#### EasySci-ATAC

Cycling MCF7 cells were barcoded with six separately loaded Tn5 N5/N7 combinations as part of the EasySci-ATAC protocol[25], which requires custom loading of adapters. We profiled the chromatin accessibility of 2’672 single cells passing quality control filters for read depth (>1’000) and overall alignment (>80% reads mapped). The majority of cells had TSS enrichment scores >10 with coverage between 5’000-10’000 fragments (Fig. 2G).

The SnapATAC2[26] framework was used for analysis, applying the Leiden community detection algorithm to identify subclusters (see Methods). As the underlying data is derived from individual cells taken from a single flask of cycling MCF7 breast cancer cells, we expected to find minimal separation of clusters, consistent with what we observed (Fig. 2H). Using Leiden annotations as a means to partition cells, we generated pseudobulk libraries to compare standard quality control metrics with bulk libraries; we observed each Leiden-based pseudobulk dataset had the same characteristic fragment length distributions and TSS enrichment profiles as seen for bulk ATAC libraries (Fig. 2E,F). Cells profiled across eight different PCR libraries distributed evenly across the landscape, revealing no library-based bias. Although generally cell-specific chromatin accessibility was not variable enough to drive distinct clustering, imputed gene expression values based on gene body accessibility exhibited distinct patterning among select cell cycle progression genes (Cyclin D1, *CCND1*), (Cyclin E2, *CCNE2*), (Chromatin Licensing and DNA Replication Factor 1, *CDT1*, which was anti-correlated with *CDKN2A*, the gene encoding the endogenous CDK inhibitor p16 (Fig. 2H).

## Discussion

OpenTn5 offers a robust and scalable solution for the production of Tn5 transposase, addressing key challenges in enzyme expression, purification, and storage. Our optimized, open-source protocol synthesizes the existing knowledge base into a robust protocol that yields multi-milligram quantities of soluble, long-term stable 10His-pG-Tn5^E54K, L372P^ per liter of *E. coli* culture. The produced Tn5 demonstrates functional equivalence to commonly used commercial enzymes, applied both in techniques which use antibody recruitment e.g. CUT&Tag, and those which do not, such as ATAC-seq. We demonstrated pG-Tn5 can be readily applied to sensitive emerging applications that require custom adapter loading, such as EasySci-ATAC[25], underlining its versatility. We hope that the OpenTn5 protocol provides a robust foundation for facilitating further developments of tagmentation, such as leveraging the development of novel Tn5 mutants, functionalization with custom protein tags, as well as DNA adapter design for emerging DNA sequencing platforms.

## Supporting information

OpenTn5 Protocol Reagents Table

OpenTn5 Protocol Buffer Table

OpenTn5 Protocol 10His-pG-Tn5(E54K L372P) Purification and Lambda Tagmentation Assay

OpenTn5 pETv2 10His-pG-Tn5 (E54K, L372P) DNA Sequence as GenBank

OpenTn5 pETv2 10His-pG-Tn5 (E54K, L372P) SnapGene DNA Sequence

## Contributions

J.S.: conceived the project, plasmid construct design, protocol development and optimization, data analysis and presentation, manuscript writing and editing with input from all authors.

L.J.A.: carried out ATAC-seq experiments, analysis, data visualization and curation.

J.L.Y.: carried out CUT&Tag experiments, analysis, data visualization and curation.

J.M.R.: carried out EasySci-ATAC experiments, analysis, data visualization, curation and provided mentorship.

D.A.O.: mentorship, protocol development and optimization.

H.A.K.: mentorship, protocol development and optimization.

V.I.R.: mentorship, funding, project supervision, manuscript writing and editing.

## Acknowledgments

J.S. acknowledges support from Boehringer Ingelheim Fonds PhD Fellowship. V.I.R. acknowledges support from NIH Common Fund (1DP2GM150021-01), The Rita Allen Foundation (Scholar Award), and Irma T. Hischl/Monique Weill-Caulier Trust (Career Science Award). J. L. Y. acknowledges support from a NSERC PGS D scholarship. L.J.A acknowledges support from the NSF Graduate Research Fellowship Program (1946429). J. M. R. was supported by the American Cancer Society – Fairfield Research Council– Postdoctoral Fellowship, PF-23-1034949-01-CCB, https://doi.org/10.53354/ ACS.PF-23-1034949-01-CCB.pc.gr.168157 H.A.K. was supported by the Uehara Memorial Foundation Research Fellowship and the Japan Society for the Promotion of Science Overseas Research Fellowships.

We acknowledge support from the Rockefeller University Structural Biology Resource Center RRID: SCR_017732. We acknowledge technical support from Rockefeller University high performance computing. We thank Wei Zhu for technical assistance. We thank Ziyu Lu and Junyue Cao for providing EasySci- ATAC adapters and protocol advice. We thank Oscar Alejandro Perez for graphic illustration and data visualization advice. We thank all members of the Risca laboratory, Yasuhiro Arimura and Pierre-Jacques Hamard for insightful discussions and advice. We thank Ariana Brenner Clerkin for manuscript formatting advice.

## Methods

### Expression, Purification, and Storage of 10His-pG-Tn5^E54K L372P^

VR124 pETv2-10His-pG-Tn5 (E54K, L372P) *E. coli* strain (Addgene #198468) glycerol stock was streaked onto LB agar plates containing 50 µg/mL kanamycin and incubated overnight at 37 °C. A single colony was picked and inoculated into 25 mL TB media supplemented with kanamycin (50 µg/mL), and the culture was grown overnight at 37 °C with shaking at 300-350 rpm. The next morning, the culture was scaled up to 500 mL TB media in a 2 L double-baffled flask, supplemented with kanamycin (50 µg/mL) and antifoam-204. The culture was grown at 37 °C until OD600 reached approximately 1.0. IPTG was then added to a final concentration of 0.6 mM to induce protein expression, and the culture was incubated overnight at 18 °C with shaking. Cells were harvested by centrifugation at 8,000 xg for 30 minutes at 4 °C, subsequently the pellets were washed by resuspension with ice-cold DPBS, pelleted and frozen at −80 °C. For lysis, the frozen cell pellets were resuspended on ice in ice-cold lysis buffer LysEQ (20 mM HEPES pH 7.8, 800 mM NaCl, 20 mM Imidazole, 10% Glycerol, 1 mM EDTA, 2 mM TCEP-HCl, 1x EDTA-free cOmplete protease inhibitor) and lysed using sonication in a salt ice bath. The lysate was clarified by centrifugation at 20,000 xg for 35 minutes at 3 °C and filtered through a 0.45 µm PES syringe filter. The cleared lysate was applied to a HisTrap HP 5 mL column (Cytiva) pre-equilibrated with buffer LysEQ, and the column was washed with buffer WashB1 (20 mM HEPES pH 7.8, 800 mM NaCl, 30 mM Imidazole, 10% Glycerol, 1 mM EDTA, 2 mM TCEP-HCl) and buffer WashB2 (20 mM HEPES pH 7.8, 800 mM NaCl, 45 mM Imidazole, 10% Glycerol, 1 mM EDTA, 2 mM TCEP-HCl) before elution with elution buffer EluB (20 mM HEPES pH 7.8, 800 mM NaCl, 400 mM Imidazole, 10% Glycerol, 1 mM EDTA, 2 mM TCEP-HCl). The eluted protein fractions were analyzed by SDS-PAGE, and the fractions containing 10His-pG-Tn5 were pooled and dialyzed overnight against SECB buffer (50 mM Tris-HCl pH 7.5, 800 mM NaCl, 10% glycerol, 0.2 mM EDTA, 2 mM DTT) in preparation for size exclusion chromatography (SEC). The following morning, the dialyzed protein was filtered through a 0.45 µm PES syringe filter. SEC was carried out on a ÄKTA pure™ FPLC system using a HiLoad 16/600 Superdex 200 pg column (Cytiva). 5 mL of filtered dialyzed protein was injected manually onto the pre-equilibrated column, followed by chromatography with 1.5 column volumes of buffer SECB at a flow rate of 1 mL/min, collecting 0.8 mL fractions from 40 mL to 110 mL elution volumes in a 96 deep-well plate. Peak fractions were analyzed by SDS-PAGE, fractions corresponding to the pure homodimer peak at 71.6 mL were pooled, and concentrated using a using a 30 MWCO Amicon Ultra concentrator to 2 mg/mL The concentrated pG-Tn5 homodimer was gently mixed with an equal volume of 100% UltraPure glycerol to a final concentration of 55% glycerol, aliquoted (1 mL) into 1.5 mL protein LoBind® tubes (Eppendorf), passively frozen and stored at −20 °C in an enzyme cooler block.

### 10His-pG-Tn5^E54K L372P^ *in vitro* tagmentation activity characterization

Nextera Mosaic End Double Stranded (MEDS) adapters were used, ordered as desalted annealed duplexes. The activity was dependent on the proper preparation of the MEDS adapters. Lambda phage genomic DNA (λ gDNA) from NEB was used as the substrate for tagmentation, and the activity was performed in the final 1xTD tagmentation buffer. To prepare the transposomes, 10 µL of pG-Tn5 stock was mixed with 10 µL of MEDS adapters and incubated at 25 °C for 10 minutes. Unless otherwise specified, the adapters were diluted to 20 µM in terms of MEDS sequence common to both N5 and N7 Nextera adapters. The tagmentation reaction was performed by mixing the transposomes with λ gDNA and incubating at 55 °C for 10 minutes. The reaction was quenched with 4% SDS, and the samples were analyzed using a genomic DNA Screentape on an Agilent Tapestation instrument or a 1% agarose EX E-Gel. One Tagment Unit (TU) of Tn5 was defined as the amount needed to uniformly tagment 125 ng of λ gDNA to <250 bp size.

### Cell culture

K562 cells were cultured in Iscove’s Modified Dulbecco’s Medium with 10% Fetal Bovine Serum and 1% PenicillinStreptomycin.

MCF-7 cells were cultured in in 75 cm^2^ flasks DMEM/F-12 (ThermoFisher 10565018) supplemented with GlutaMAX (Gibco 10565018), 10% Fetal Bovine Serum and 1% Penicillin-Streptomycin at 37 °C and 5% CO_2_. Cells were subcultured at a 1:3 ratio every 3 days. Prior to their use, MCF7 cell identity was verified by STR profiling through LabCorp Cell Line Authentication service.

### Library Preparation

#### pG-Tn5 adapter loading

20 µM Nextera sequencing adapters were in a 1:1 volume with pG-Tn5, mixed gently by pipetting, incubated at 23 °C for 10 minutes, and used same-day.

#### CUT&Tag

CUT&Tag on MCF7 cells was performed by following the protocol from Kaya-Okur et al, Nature Protoc 2020[27]. MCF7 cells were briefly centrifuged followed by incubation with Nuclear Extraction buffer on ice for 10 minutes. After resuspension in Wash Buffer, nuclei were slow frozen in cryogenic vials with 10% DMSO for storage at −80 °C before resuming the rest of the protocol. Approximately 100,000 cells were used per sample for overnight incubation with primary antibody (Tri-Methyl-Histone H3 (Lys27) (C36B11) Rabbit mAb, 9733T, Cell Signaling Technology 1:50, Anti-Histone H3 (acetyl K27) antibody - ChIP Grade (ab4729), Abcam 1:100). Secondary antibody (Guinea Pig anti-Rabbit IgG (Heavy & Light Chain) antibody, Guinea Pig, ABIN101961 1:100) was incubated for 30 minutes at room temperature followed by addition of commercial pAG-Tn5 (EpiCypher SKU: 15-1017) or inhouse pG-Tn5 preloaded with adapters. Phenol chloroform DNA extraction was performed using Qiagen’s Maxtract High Density phase-lock tubes. Libraries were PCR amplified for 12 cycles with Illumina i5 or i7 barcoded primers and Ampure XP beads were used for post-PCR cleanup.

#### ATAC-seq

ATAC assay was performed in duplicate or triplicate on 50,000 cells per technical replicate following Omni-ATAC protocol with the following modifications[11]. Prior to tagmentation, cells were treated in culture medium with DNase (Worthington cat# LS002007) at a final concentration of 200 U/mL at 37 °C for 30 mins, then washed twice with PBS at 500g, RT. For Illumina enzyme tagmented material, 2.5 µL of Illumina TDE1 enzyme was used. K562 transposome titration experiment: pG-Tn5 tagmentation was titrated with the following amounts of loaded transposome: 1.0, 2.5, 5.0, 7.0 µL. 7 cycles of PCR were performed during library preparation for K562 experiment. MCF7: 8 cycles of PCR were performed during library preparation of MCF7 experiment. Prior to sequencing, fragment length distribution and DNA concentration were determined via Tapestation and Qubit respectively.

#### EasySci-ATAC

For harvesting, flasks were washed with ice cold PBS then cells were dissociated with TrypLE Express enzyme. Nuclei were extracted and single-cell indexing was performed following the EasySci-ATAC protocol, as described in Supplemental protocol 2 of Sziraki, et al., 2023[25]. Briefly, cells were resuspended in 1 mL Nuclei Isolation Buffer (NIB) + 0.1% IGEPAL CA-630 and filtered through a 40 µm filter. Nuclei were stained with Trypan Blue and counted with the EVE Automated Cell Counter; the final concentration was adjusted to 1’000 nuclei/µL using NIB. For each barcode combination, 5’000 nuclei were mixed with 4 µL of barcoded pG-Tn5 and tagmentation was carried out in a thermomixer at 55 °C for 10 minutes with gentle shaking at 350 RPM. Following tagmentation, nuclei were put directly on ice to prevent over tagmentation, followed by addition of a stop buffer consisting of spermidine-EDTA. Nuclei were then pooled and redistributed to 96-wells to allow multiplexing via barcoded ligation, followed by a third round of pooling and redistribution to a final 96-well reaction for barcoding via PCR amplification and library preparation. Libraries underwent size selection via E-gel prior to sequencing. All libraries were sequenced to 0.5-10M reads on the NextSeq 1000 with 58,8,8,60 sequencing parameters.

### Custom loading of Tn5 with combinatorial barcodes for EasySci-ATAC

For uniquely barcoded tagmentation reactions, two N5 barcoded oligo duplexes were mixed in combination with three N7 barcoded oligo duplexes. Per barcode combination: 1 µL of 10 µM N5 duplex, 1 µL of 10 µM N7 duplex, and 2 µL of pG-Tn5 (Lot 2) were mixed and incubated for 15 minutes at 25 °C. Loaded pG-Tn5 was kept on ice until nuclei were ready for tagmentation.

### Data Analysis for bulk CUT&Tag and ATAC-seq

#### Adapter trimming

Adapters were trimmed from pair-end FASTQ files using a custom script adapted from Corces et al. 2017[11] (https://github.com/riscalab/pipeSeq/blob/master/scripts/pyadapter_trimP3V2.py) (https://github.com/riscalab/pipeSeq/blob/master/scripts/pyadapter_trimP3V2.py), then FASTQC and FASTQC Screen performed per-base sequence content QC and screened for mycoplasma genome representation.

### Alignment to hg38

#### CUT&Tag

Reads were aligned to hg38 using bowtie2[28] using the following parameters--end-to-end --very-sensitive --no-mixed --no-discordant --phred33 -I 10 -X 700 for mapping of inserts 10-700 bp in length.

#### ATAC-seq

Reads were aligned using -X 2000 parameters on bowtie2. Reads were shifted to account for Tn5 insertion bias using deepTools[29] alignmentSieve command with —ATACshift parameter set.

For both CUT&Tag and ATAC-seq data, duplicate reads, reads mapping to the mitochondrial genome and blacklisted regions were removed using picardtools[30] to generate the final bam file. Bam files were then sorted using picardtools. Libraries sequenced across multiple flow cells were merged as bams, then sorted using picardtools.

### Peak Calling

#### CUT&Tag

Using bedtools, bam files were converted to bedpe files which were then converted to bedgraph format. Bedgraph files were used as input into SEACR peak caller[31], where peaks were called with parameters specified to non-normalized and stringent threshold set to 0.01. Afterwards, a master peak set between technical replicates was created keeping only peaks that overlapped with each other between technical replicates and merging overlapping peaks into a single peak. A master non-redundant peak set was created by merging overlapping peaks of peaksets from technical replicates of different Tn5 conditions (Epicypher’s pAG-Tn5 vs in house pG-Tn5) using GenomicRanges package[32] in R.

#### ATAC-seq

Peaks were called from bam files with MACS2[33] using the following parameters -B --SPMR -- nomodel --shift -37 --extsize 73 --nolambda --keep-dup all --call-summits --slocal 10000. Next, a non-redundent master peakset was made by intersecting technical replicate narrowPeaks in bedtools, then merging intersected narrowPeak files in bedtools for each cell type. Cell-type masterpeak sets were merged in bedtools[34] to create a non-redundant peakset across all ATAC samples.

### Sample Correlation

Fragment counts for each replicate under the masterpeak set were counted using chromVAR’s getCounts function[35]. For K562 transposome dilution analysis, fragment counts under 1 kb hg38 bins were counted using chromVAR’s getCounts function for each replicate[35]. The resultant fragment count matrix was then used as input into DESeq2[36]v1.26.0 and sequencing depth was normalized across samples using the estimateSizeFactors function.For visualization purposes, the counts matrix underwent a “regularized log” transformation using the rlog function from DESeq2. Next, the distance measure between pairwise samples was calculated. 1 – distance measure was used as input to calculate the Spearman correlation coefficient between each sample pair. Custom python matplotlib code was used to plot the sample correlation matrix.

### *E. coli* transposome contamination analysis

Reads were aligned using bowtie2 to the *E. coli* genome from ATAC-seq bam files. Number of *E. coli* mapped reads/Total mapped reads were calculated and plotted.

### Genome coverage visualization

Technical replicates were combined into a single bam file using samtools merge command. Bigwig files were generated using deepTools’ bamCoverage command with the following parameters to normalize:

Coverage to the hg38 genome size CUT&Tag: --binSize 5 --normalizeUsing RPGC --effectiveGenomeSize 2913022398 --extendReads.

Coverage to the hg38 genome size ATAC-seq: --binSize 1 --normalizeUsing RPGC --effectiveGenomeSize 2913022398 --extendReads

### ATAC-seq Coverage Plots

#### Plotting CTCF Insertions

Merged ATAC-seq bam files from technical replicates were randomly down-sampled to 3M reads using samtools for read-depth normalized comparisons. Homo sapiens CTCF PWM from the HocomocoV10 database[37] were used to plot insertions +/-100bp around the motif using the regionPlot function from soGGI R package[38].

#### TSS Enrichmen

TSS enrichment profile of insertions were produced as previously described[11] using hg38 TSS annotation from gencode. Insertions were counted in 1bp bins +/-2’000 bp surrounding annotated TSSs. Bins were then normalized by dividing by the average of the first 200 bp bins from each flank: https://github.com/riscalab/pipeSeq/blob/master/scripts/pyMakeVplot_css_v01.py https://github.com/riscalab/pipeSeq/blob/master/scripts/pyMakeVplot_css_v01.py

#### Coverage CTCF

IDR-thresholded CTCF narrow peak ChIP-Seq data in MCF7 cell line was downloaded from ENCODE accession number ENCFF278FNP https://www.encodeproject.org/files/ENCFF278FNP/, https://www.encodeproject.org/files/ENCFF278FNP/. Total coverage around ChIP CTCF peaks was calculated using deepTools computeMatrix function on RPGC normalized bigwigs (described above) in reference-point mode with 1 kb flanking regions. The matrix was plotted using deepTools plotHeatmap function with bilinear interpolation.

### Computational procedures for processing EasySci-ATAC libraries

Libraries were demultiplexed using custom scripts to assign base calls to individual cells based on indices from PCR, ligation, and Tn5 combinatorial barcoding. The SnapATAC2 analysis framework was followed to generate the TSS enrichment score vs. unique fragments plot, perform doublet removal, dimensional reduction, and Leiden algorithm derived clustering analysis[26]. Gene body accessibility was counted using Gencode v38 basic gene annotation, and subsequently used for visualizing gene accessibility as a proxy for expression after applying the MAGIC algorithm for imputation and data smoothing.

## Supplementary Files

1. OpenTn5 Protocol 10His-pG-Tn5^E54K, L372P^ Purification and Lambda Tagmentation Assay
2. OpenTn5 Protocol Buffer Table
3. OpenTn5 Protocol Reagents Table
4. Supplementary Table 1 (Sequencing QC metrics)
5. Plasmid sequence as .genbank and .dna file
6. Raw and Processed Sequencing Data (GEO)

**Figure S1:**
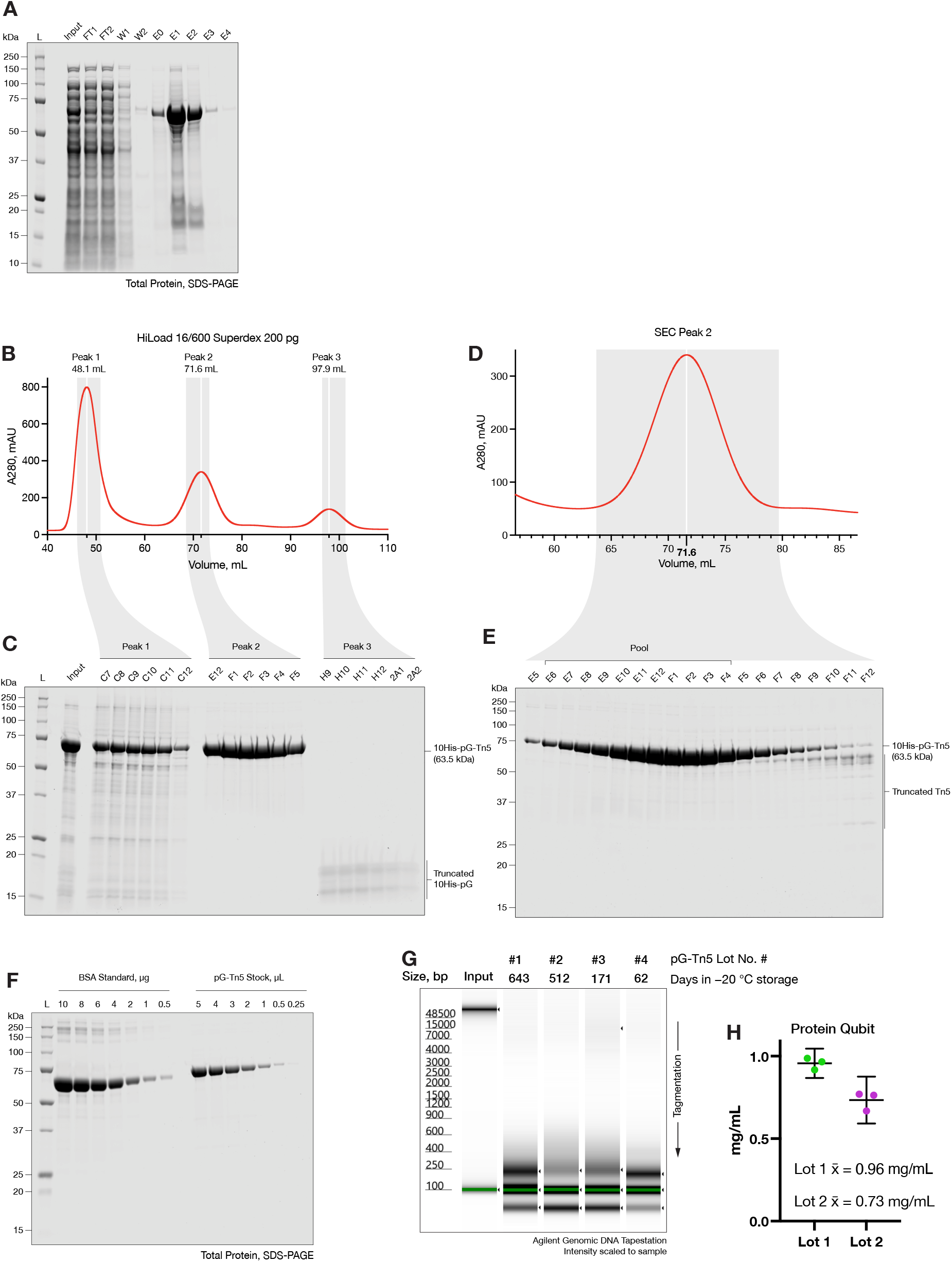
Overview of pG-Tn5 purification, validation of stock concentration, stability testing. **A**. Profiling of total protein content from His-tag affinity purification fractions, InstantBlue stained SDS-PAGE. **B**. A280 profile of 10His-pG-Tn5 size exclusion chromatography (SEC) carried out on pooled His-tag affinity eluate fractions from panel A above, on HiLoad 16/600 Superdex 200 pg column using an ÄKTA FPLC instrument. **C**. SDS-PAGE analysis of highlighted SEC peak fractions corresponding to 10His-pG-Tn5 containing heterogeneous aggregates (peak 1), 10His-pG-Tn5 homodimer (peak 2) and truncated 10His-pG tag (peak 3). **D**. Detail of SEC peak 2, from the same purification run as panel B. **E**. SDS-PAGE of fractions corresponding to the highlighted region of SEC peak 2, showing contaminating 10His-pG-Tn5:Tn5 heterodimers containing truncated Tn5, which migrate slower than full length 10His-pG-Tn5 homodimer. SEC fractions containing pure, full-length 10His-pG-Tn5 homodimer chosen for pooling to generate stock are highlighted. **F**. Validation of protein concentration of 1 mg/mL 10His-pG-Tn5 final stock by dilution series against a BSA standard, analyzed by titration on SDS-PAGE with InstantBlue total protein staining. **G**. Assaying 10His-pG-Tn5 stock tagmentation activity after indicated time stored at −20 °C, showing negligible loss of λ gDNA tagmentation activity at 21 months. Equal volumes of stock were used to assemble 20 µM adapter loaded transposomes. **H**. Quantification of protein concentration of pG-Tn5 stock lots 1 and 2, demonstrating that the tagmented DNA product size is largely attributable to the variation in stock protein concentration.

**Figure S2:**
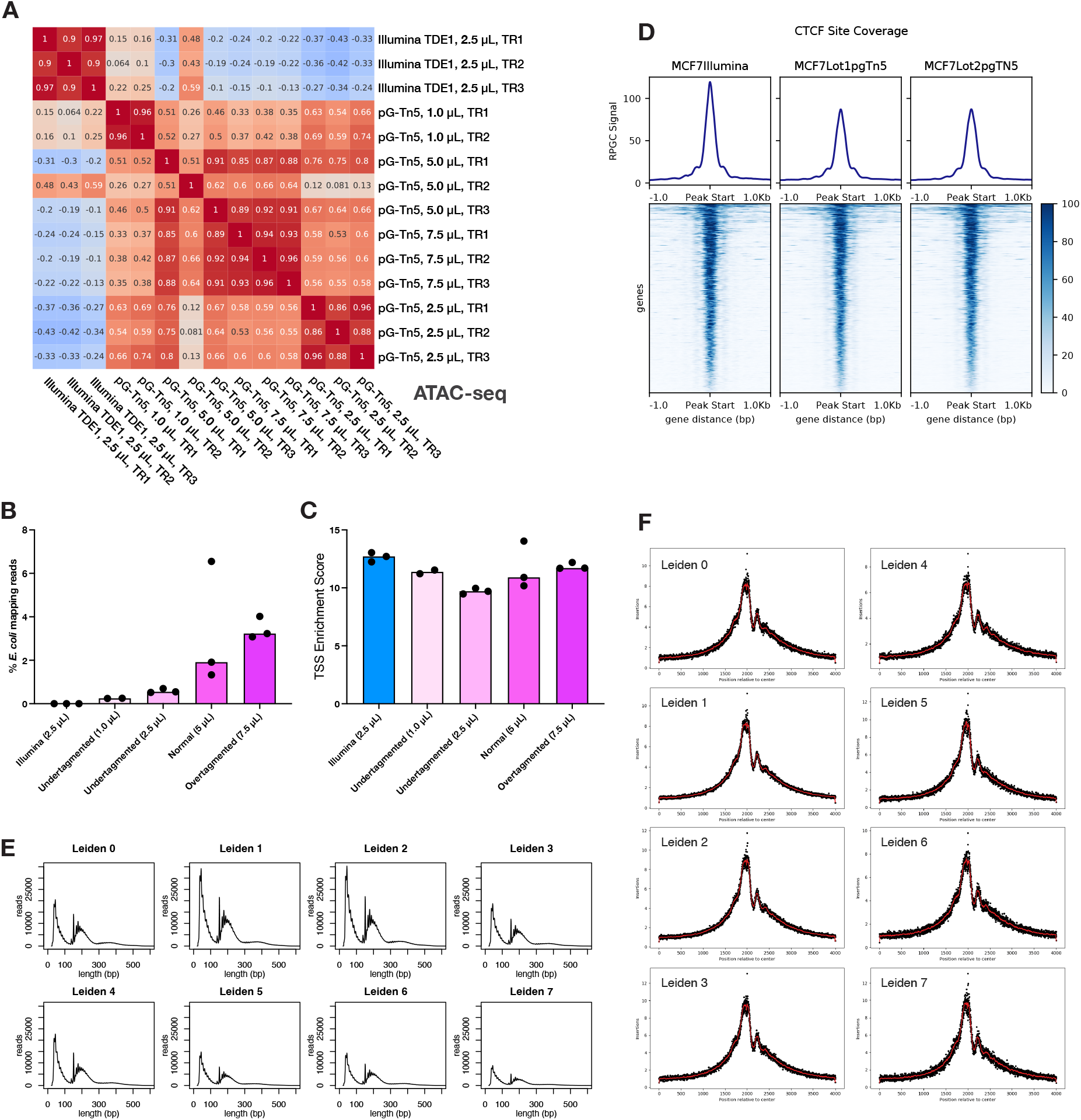
Validating pG-Tn5 in bulk and single-cell ATAC-seq performance using differing transposome volumes, bulk ATAC-seq coverage over individual CTCF sites, and EasySci-ATAC data quality metrics. **A**. Spearman rank correlation of ATAC-seq libraries generated in human K562 cells using different volumes of pG-Tn5 transposome, as compared to standard OmniATAC-seq protocol 2.50 µL volume of Illumina TDE1, calculated over 1 kb bins genome-wide. **B**. Relationship between pG-Tn5 transposome volume used per ATAC-seq reaction and % library reads aligning to *E. coli* genome, originating from carryover during purification, data from same experiment as in panel A above. **C**. Transcription start site (TSS) enrichment score of ATAC-seq libraries generated with Illumina TDE1 or differing pG-Tn5 transposome volumes, data from the same experiment as in panel A above. **D**. Heatmap of ATAC-seq signal coverage of individual CTCF-occupied sites in human MCF7 cells, as in Figure 1D-F. **E**. Fragment length distribution (FLD) plots of EasySci-ATAC individual Leiden annotations. **F**. Transcription start site (TSS) enrichment score plots of EasySci-ATAC individual Leiden annotations.

