## Supplementary material for "OpenTn5: Open-Source Resource for Robust and Scalable Tn5 Transposase Purification and Characterization": OpenTn5 Protocol 10His-pG-Tn5(E54K L372P) Purification and Lambda Tagmentation Assay

### 10His-pG-Tn5$^{E54K, L372P}$ OpenTn5 Transposase Purification and Characterization Protocol

**Reagents and Consumables**

*This table is also provided as a supplementary file "OpenTn5 Protocol Reagents Table.xlsx"*

| Reagent / Consumable | Manufacturer | Catalog # |
|---|---|---|
| 1 Liter IDTE pH 7.5 (10 mM Tris, 0.1 mM EDTA) 1X TE Solution | IDT | 11-05-01-05 |
| Amicon® Ultra Centrifugal Filter, 30 kDa MWCO, sample volume 15 mL | Millipore | UFC903008 |
| Antifoam 204 | Sigma-Aldrich | A6426-100G |
| Any kD™ Criterion™ TGX™ Precast Midi Protein Gel, 26-well, 15 µL | Bio-Rad | 5671125 |
| Any kD™ Mini-PROTEAN® TGX™ Precast Protein Gels, 15-well, 15 µL | Bio-Rad | 4569036 |
| cOmplete™, EDTA-free Protease Inhibitor Cocktail | Sigma-Aldrich | 11873580001 |
| Cytiva Whatman™ Uniflo Syringe Filters, 0.45 µm PES | Cytiva | 09-928-063 |
| DTT (Dithiothreitol) (> 99% pure) Protease free | Goldbio | DTT25 |
| E-Gel™ 1 Kb Plus Express DNA Ladder | Invitrogen | 10488091 |
| E-Gel™ EX Agarose Gels, 1% | Invitrogen | G402021 |
| Fisher BioReagents™ Terrific Broth | Fisher Scientific | BP2468-500 |
| Gibco™ DPBS, no calcium, no magnesium | Gibco | 14-190-144 |
| Glycerol (Molecular Biology), Fisher BioReagents™ | Fisher Scientific | BP229-4 |
| HEPES, Free Acid | Goldbio | H-400-2 |
| HisTrap HP, 1 x 5 ml | Cytiva | 17524801 |
| Imidazole, Recrystallized | Goldbio | I-902-100 |
| InstantBlue® Coomassie Protein Stain | Abcam | ab119211 |
| Invitrogen Qubit Assay Tubes | Invitrogen | Q32856 |
| Invitrogen Qubit Protein Assay, 12.5 µg/mL to 5 mg/mL range | Invitrogen | Q33211 |

| | | |
|---|---|---|
| IPTG | Goldbio | I2481C25 |
| Kanamycin monosulfate, ≥750 U/mg, USP Grade, Powder | Goldbio | K-120-25 |
| Lambda DNA, New England Biolabs, Lambda DNA, Size=250 µg, 500 µg/mL | NEB | N3011S |
| LB agar, Miller | Sigma-Aldrich | L3027-1KG |
| MgCl2 (1 M) | Invitrogen | AM9530G |
| Millex-GP Syringe Filter Unit, 0.22 µm, polyethersulfone, 33 mm, gamma sterilized | Millipore | SLGPR33RS |
| N,N-Dimethylformamide (DMF), anhydrous, 99.8% | Thermo Scientific Chemicals | 043997.K2 |
| Nalgene™ Rapid-Flow™ 500mL Bottle Top Filter, PES filter material, 0.2µm, 75mm membrane, Sterile, 45mm neck | Nalgene | 595-4520 |
| Pierce BCA Protein Assay Kit with Dilution-Free BSA Protein Standards | Pierce | A55864 |
| Precision Plus Protein™ Unstained Protein Standards | Bio-Rad | 1610363 |
| Protein LoBind® Tubes, 1.5 mL, snap cap, PCR clean | Eppendorf | 22431081 |
| Protein LoBind® Tubes, 15 mL, conical tubes, PCR clean | Eppendorf | 30122216 |
| Protein LoBind® Tubes, 5.0 mL, snap cap, PCR clean | Eppendorf | 30108302 |
| Protein LoBind® Tubes, 50 mL, conical tubes, PCR clean | Eppendorf | 30122240 |
| SDS, 20% Solution, RNase-free | Invitrogen | AM9820 |
| Slide-A-Lyzer™ G3 Dialysis Cassettes, Irradiated 10K MWCO, 15 mL sample capacity | Thermo Scientific | A52982 |
| Sodium Chloride | Supelco | SX0420-5 |
| TCEP HCl | Goldbio | TCEP5 |
| UltraPure™ 0.5M EDTA, pH 8.0 | Invitrogen | 15575020 |
| UltraPure™ 1 M Tris-HCl Buffer, pH 7.5 | Invitrogen | 15567027 |
| UltraPure™ DNase/RNase-Free Distilled Water | Invitrogen | 10977035 |
| UltraPure™ Glycerol | Invitrogen | 15514011 |

#### Equipment

*This list is non exhaustive, but includes key specialized equipment required to follow the OpenTn5 protocol with minimal modifications. This table is also provided as a supplementary file “OpenTn5 Protocol Reagents Table.xlsx”*

| Equipment | Manufacturer | Catalog # |
|---|---|---|
| 4150 TapeStation System | Agilent | G2992AA |
| Flask, Shake, 2000mL, 6 Baffles, GL-45 Thread, Vented Cap | Chemglass | CLS-2052-10 |
| Q125 Sonicator | Qsonica | Q125-110 |
| Criterion™ Cell, Vertical midi-format electrophoresis cell | Bio-Rad | 1656001 |
| Mini-PROTEAN® Tetra Vertical Electrophoresis Cell for Mini Precast Gels, 4-gel | Bio-Rad | 1658004 |
| E-Gel™ Power Snap Electrophoresis Device | Invitrogen | G8100 |
| Qubit™ 4 Fluorometer, with WiFi | Invitrogen | Q33238 |

**General equipment**

Chilled centrifuge capable of >8'000 xg at 4 °C

Gel imager

MilliQ water supply

Peristaltic pump setup

FPLC system with A280/A260 spectrophotometer

Chilled shaker incubator capable of chilling to 18 °C

Tabletop spectrophotometer, Nanodrop or equivalent

#### Buffer Compositions

| | |
|---|---|
| **1x LysEQ** | 20 mM HEPES pH 7.8, 800 mM NaCl, 20 mM Imidazole, 10% Glycerol, 1 mM EDTA, 2 mM TCEP-HCl |
| **1x WashB1** | 20 mM HEPES pH 7.8, 800 mM NaCl, 30 mM Imidazole, 10% Glycerol, 1 mM EDTA, 2 mM TCEP-HCl |
| **1x WashB2** | 20 mM HEPES pH 7.8, 800 mM NaCl, 45 mM Imidazole, 10% Glycerol, 1 mM EDTA, 2 mM TCEP-HCl |
| **1x EluB** | 20 mM HEPES pH 7.8, 800 mM NaCl, 400 mM Imidazole, 10% Glycerol, 1 mM EDTA, 2 mM TCEP-HCl |
| **1x SECB** | 50 mM Tris-HCl pH 7.5, 800 mM NaCl, 10% glycerol, 0.2 mM EDTA, 2 mM DTT |
| **2x Tagmentation Buffer** | 20 mM Tris-HCl pH 7.5, 10 mM MgCl2, 20% DMF |

---

#### Buffer Recipes

*Detailed buffer recipe tables are provided as a supplementary file "OpenTn5 Protocol Buffer Table.xlsx"*

---

#### OpenTn5 Protocol

##### Section A: 10His-pG-Tn5$^{E54K, L372P}$ Expression, Purification and Storage

**Day 1**
**Glycerol stock streak-out**
It is best practice to begin protein production from a freshly streaked out *E. coli* glycerol stock.

1. Streak out VR124 pETv2-10His-pG-Tn5$^{E54K, L372P}$ (Addgene #198468) glycerol stock on an LB agar plate containing 50 µg/mL kanamycin, place plate in a 37 °C incubator overnight.

   Note: When preparing a batch of fresh LB agar kanamycin plates, verify the potency of the antibiotic by streaking out non-kanamycin resistant *E. coli*, for example by plating an aliquot of competent cells. Repeat this test if using LB agar plates that you are unsure the freshness of.

**Day 2**
**Starter culture**
Start in the evening.

2. Pick a single colony from the overnight-incubated LB agar kanamycin plate and inoculate into a 250 mL culture flask containing 25 mL of TB media supplemented with 25 µL of 1000x 50 mg/mL kanamycin stock, to give a final concentration of 50 µg/mL kanamycin. Shake at 300–350 rpm at 37 °C overnight.

Note: Good culture aeration is essential for healthy *E. coli* growth, especially when using a rich media such as TB. To achieve this in small scale cultures, do not exceed the media volume comprising 25% of culture flask volume and shake at speeds > 250 rpm.

**Day 3**
**Culture scale-up and induction**
Start first thing in the morning. Pre-cool second shaker incubator to 18 °C.

**3.** To a 2 L double-baffled culture flask, add 500 mL of TB media, supplement with 500 µL of 1000x 50 mg/mL kanamycin stock and 30 µL of antifoam-204.

Note: Good culture aeration is critical for reliable, high-yield protein expression in *E. coli*. To achieve this in large cultures we use double-baffled 2 L flasks and fill them with no more than 500 mL media. If baffled flasks are not available, culture in multiple flasks with less media in each, for example 4x125 mL cultures in 2 L flasks. Additionally, we supplement media with antifoam-204 which reduces foam accumulation on the surface of the culture, which otherwise limits oxygenation of the culture. Higher shaking speed of 300-350 rpm also help with culture aeration.

**4.** Add 5 mL of overnight VR124 starter culture to 500 mL of supplemented TB media to give a 1:100 inoculation. Shake at 300–350 rpm at 37 °C until culture reaches $OD_{600}$ ≈ 1.00, which should take about 2 hours. Begin checking culture $OD_{600}$ as soon as 1-hour post-inoculation.

Note: We routinely measure culture $OD_{600}$ using a Nanodrop spectrophotometer, measuring 2 µL of undiluted culture directly. We have found this to be a sufficiently accurate and reliable alternative to cuvette measurements.

**5.** Once $OD_{600}$ ≈ 1.00, place culture on ice and take a 3 mL aliquot of culture to a 15 mL culture tube to serve as the uninduced control. Place uninduced control side-culture on ice.

**6.** Add 600 µL of 0.5 M IPTG stock to rest of the ice-chilled culture to give a final 0.6 mM IPTG concentration. Swirl the flask by hand immediately upon IPTG addition and place the 2 L flask and the uninduced 3 mL side-culture into a shaker pre-cooled to 18 °C. Shake at 300–350 rpm at 18 °C overnight for 18–24 hours.

Note: Pre-chilling the culture on ice is important since protein expression induction at lower temperatures promotes protein solubility. However, prolonged cooling of *E. coli* on ice in absence of shaking can cause cell death and should be avoided. Therefore, limit the time the cultures spend on ice, 10 minutes is usually sufficient to cool the culture to < 18 °C.

During day 3 you can prepare all buffers required for day 4, except for adding the cOmplete protease inhibitor EDTA-free tablet to the lysis buffer. Chill all buffers and DPBS to 4 °C.

**Day 4**
**Culture harvest, lysis, affinity purification and dialysis**

Start first thing in the morning. Pre-cool centrifuges you will use to 0–4 °C.

**7.** After 18–24 hours of induction, measure the $OD_{600}$ of both the main induced culture, as well as the uninduced side-culture. After 20 h of induction typical $OD_{600} \approx 7.50$, whilst the uninduced side-culture $OD_{600} \approx 10.00$.

**8.** Take small, 200 µL aliquots of the induced and uninduced cultures for validating 10His-pG-Tn5 overexpression by SDS-PAGE.

**9.** Weigh the empty centrifuge bottles you will be spinning the cultures in to later determine the wet pellet weight.

**10.** Pellet the induced culture in appropriate centrifuge bottles at 8'000 xg for 30 minutes at 0–4 °C

**11.** Remove and discard all the media supernatant, which should be clear, and weigh the centrifuge bottles containing the wet pellets. 500 mL of induced culture at $OD_{600} \approx 7.50$ should give ≈ 7 g wet pellet.

**12.** Wash the pellets from 500 mL culture by resuspending them by pipetting in 30 mL of ice-cold DPBS, transfer to a 50 mL conical tube and spin at 4'300 xg for 15 minutes at 0–4 °C.

**13.** Carefully remove and discard all the DPBS supernatant, the final wet pellet volume from 500 mL culture at $OD_{600} \approx 7.50$ should be ≈ 5 mL.

**14.** Place conical tube containing the wet pellet in –80 °C freezer. Snap-freezing in liquid $N_2$ is not required.

Pause point: pellets may be safely kept at –80 °C at this point for > 4 months.

**15.** Allow pellets to freeze completely in the –80 °C freezer for > 30 minutes.

Warning: Do not skip this freeze-thaw cycle, as it helps with disrupting the *E. coli* cell walls and membranes during subsequent lysis.

**16.** Whilst the pellet is in the –80 °C freezer, perform a quick SDS-PAGE analysis of both uninduced and induce culture aliquots taken earlier.

**17.** Spin the 200 µL aliquots of uninduced and induced cultures at 5'000 xg for 10 minutes at 0–4 °C.

**18.** Remove and discard the media supernatant from both culture aliquots and resuspend the pellets in ice-cold DPBS by pipetting. Resuspend the induced sample in 200 µL DPBS. Adjust the volume of DPBS used to resuspend the uninduced pellet by the $OD_{600}$(uninduced)/ $OD_{600}$(induced) ratio. For example, 10.0/7.50 = 1.34 x 200 µL = 268 µL DPBS to resuspend 200 µL uninduced culture at $OD_{600}$ = 10.0. Adjusting the DPBS volume at this stage will normalize the cell mass concentration between the samples, making 10His-pG-Tn5 over-expression comparison in SDS-PAGE easier.

**19.** For both the uninduced and induced samples, lyse 2.00 µL of *E. coli* resuspended in DPBS directly in loading buffer, denaturing at 90 °C for 10 minutes in presence of 50 mM DTT final in loading buffer.

Note: Over-loading SDS-PAGE gels makes differential band comparison difficult, therefore consider running several dilutions of the samples alongside on the same gel.

**20.** Run denaturing SDS-PAGE mini-gel following manufacturer's instructions.

**21.** Quickly rinse the gel several times in deionized $H_2O$, stain with ≈ 30 mL InstantBlue Coomassie protein stain, rotating for >1 hour, followed by destaining in deionized $H_2O$. Bands should become visible after 15 minutes, with a clear intense 10His-pG-Tn5 band at ≈ 63.5 kDa present only in the induced sample.

Note: Absence of clear 10His-pG-Tn5 over-expression at this step requires troubleshooting of the preceding protocol steps.

**22.** Add one Roche cOmplete EDTA-free protease inhibitor tablet to a 50 mL aliquot of 1x LysEQ buffer. Make sure the tablet is fully dissolved before proceeding.

**23.** Move the conical tube containing the pellet stored at –80 °C onto ice, add 25 mL of protease inhibitor-supplemented 1x LysEQ to the pellet and begin resuspending the pellet immediately on ice, so it thaws as it is resuspended. Pipette thoroughly on ice until no visible pellet clumps remain, minimize foaming.

**24.** Prepare a salt ice bath in a 1 L glass beaker by adding ≈ 150 g NaCl, filling the beaker with ice and adding deionized water to fill the beaker about halfway. There is no need to mix.

**25.** Place the conical tube with the resuspended pellet into the salt ice bath and let pre-chill for 30 minutes prior to beginning sonication.

**26.** Place the ice bath beaker containing the conical tube into the sonicator probe enclosure, adjusting the height of the platform so that the sonicator probe is submerged 1/3 of the depth of the lysate.

**27.** Sonicate the lysate, 80% amplitude, 10 seconds ON, 10 seconds OFF, for a total of 6 minutes ON time.

Warning: During sonication the conical tube will sink as the ice melts. Monitor the submersion level of the probe throughout the sonication, adjusting the tube in the ice bath to keep the probe submerged 1/3 of the depth of the lysate. The lysate should visibly circulate in the tube during the ON periods but should not froth excessively. Excessive heating of the lysate during sonication can denature 10His-pG-Tn5, resulting in cleavage of the 10His-pG tag and subsequent loss of yield of full-length 10His-pG-Tn5 homodimer.

Note: These sonication settings are specific to our QSonica sonicator setup for ≈ 30 mL lysate from 500 mL culture at $OD_{600} \approx 7.50$. These settings will vary with your sonication conditions, so you will need to empirically test them yourself. If processing larger culture volumes it is preferable to sonicate them in batches, or empirically test sonication conditions of larger lysate volumes.

**28.** Check the viscosity of the lysate by pipetting using a P200 pipet. Sufficiently sonicated lysate should come out of the P200 tip as a stream instead of viscous droplets.

**29.** Save a 100 µL aliquot of sonicated lysate for later analysis.

**30.** Transfer the sonicated lysate to a suitable centrifuge tube, spin at 20’000 xg for 35 minutes at 3 °C.

**31.** Whilst the lysate is spinning, begin equilibration of the HisTrap HP 5 mL column cartridge.

**32.** Working in a cold room, using a peristaltic pump, wash the HisTrap column with 10 column volumes (10 CV) of cold Milli-Q $H_2O$ at 1 mL/min flowrate.

Note: We use a peristaltic pump for affinity chromatography to avoid applying lysate into the FPLC system. However, if you prefer, you may use a FPLC instrument to streamline the affinity purification steps of the protocol. Extensive equilibration with cold Milli-Q $H_2O$ is only necessary if using a freshly-opened HisTrap HP 5 mL column.

**33.** Wash HisTrap column with 10 CV of cold 1x LysEQ buffer at 1 mL/min flowrate.

**34.** Save a 100 µL aliquot of clarified lysate for later analysis.

**35.** Filter the clarified lysate through a 0.45 µm PES syringe filter, keep the filtered lysate on ice.

Note: You may need to use multiple filters if they become clogged and passing lysate becomes difficult. Alternatively, use a 50 mL conical tube-top vacuum filter unit to filter the clarified lysate.

**36.** Save a 100 µL aliquot of filtered lysate for later analysis.

**37.** Apply the filtered lysate onto the HisTrap column at a lowered flowrate of 0.8 mL/min, saving the flowthrough in a clean 50 mL conical tube. This is sample is ‘flowthrough-1’.

**38.** Save a 100 µL aliquot of flowthrough-1 for later analysis.

**39.** Apply the flowthrough-1 onto the same HisTrap column at 0.8 mL/min. Save the flowthrough in a clean 50 mL conical tube as ‘flowthrough-2’.

**40.** Wash the HisTrap column with 10 CV (50 mL) of 1x WashB1 buffer at 1 mL/min. Save the flowthrough fraction in a clean 50 mL conical tube as ‘wash-1’.

**41.** Wash the HisTrap column with 10 CV (50 mL) of 1x WashB2 buffer at 1 mL/min. Save the flowthrough fraction in a clean 50 mL conical tube as ‘wash-2’.

**42.** Elute the HisTrap column with 1 CV (5 mL) of 1x EluB buffer at 0.5 mL/min. Save the flowthrough fraction in a 15 mL conical tube as ‘elution-0’.

Note: This elution-0 fraction is largely be composed of WashB2 buffer and contains little 10His-pG-Tn5.

**43.** Incubate the HisTrap column for 30 minutes in the cold room to facilitate 10His-pG-Tn5 elution off the column.

**44.** Elute the HisTrap column with 0.5 CV (2.5 mL) of 1x EluB buffer at 0.5 mL/min. Save the flowthrough fraction in a 15 mL conical tube as ‘elution-1’.

**45.** Elute the HisTrap column with 0.5 CV (2.5 mL) of 1x EluB buffer at 0.5 mL/min. Save the flowthrough fraction in a 15 mL conical tube as ‘elution-2’.

**46.** Repeat the elution process two more times for elution-3 and elution-4 fractions, each 2.5 mL.

**47.** Analyze filtered input, flowthrough-1 (FT1), FT2, wash-1 (W1), W2, elution-0 (E0), E1, E2, E3, E4 fractions by SDS-PAGE with Coomassie staining. Load 2.0 µL of input, FT1, FT2 fractions, 4.5 µL of W1, W2 fractions, 10.0 µL of E0 fraction, 5.0 µL of E1, E2, E3, E4 fractions per well of a mini-gel.

**48.** Based on the SDS-PAGE, confirm that 10His-pG-Tn5 is depleted in the FT1 and FT2 fractions, indicating good binding to the column. The majority of 10His-pG-Tn5 should be present in E1 and E2 fractions, which are pooled together to give 5 mL of eluate pool.

Note: If significant 10His-pG-Tn5 is present in later elution fractions, you can increase the eluate pool volume. We choose to limit the volume of the eluate pool to 5 mL, since this is the maximum injection volume of the HiLoad 16/600 Superdex 200 pg column we use for size exclusion chromatography. However, if necessary, it is possible to concentrate the eluate pool post-dialysis prior to size exclusion chromatography.

**49.** Dialyze the eluate pool overnight against 2 L of pre-chilled 1x SECB buffer in a 2 L glass beaker with a magnetic stir bar in the cold room. Transfer the eluate pool to a pre-equilibrated 15 mL capacity 10K MWCO Slide-A-Lyzer™ G3 Dialysis Cassette.

Warning: Make sure to pre-equilibrate the dialysis cassette in the 1x SECB buffer for > 5 minutes, making sure both sides of the dialysis cassette are wetted in the process.

**Day 5**
**Size exclusion chromatography, concentration, and storage**

**50.** Using an ÄKTA FPLC system, equilibrate the HiLoad 16/600 Superdex 200 pg size exclusion column with 2 CV (240 mL) of pre-chilled 0.22 µm-filtered 1x SECB buffer.

Note: It is possible to use a different size exclusion column. We have successfully used both HiLoad 26/600 Superdex 200 pg as well as Superdex 200 Increase 10/300 GL columns to isolate the 10His-pG-Tn5 homodimer.

**51.** Save a 50 µL aliquot of the dialyzed elution pool.

**52.** Filter the dialyzed elution pool through a 0.45 µm PES syringe filter into a clean tube. This step removes any precipitates that may have formed during the process of dialysis and which could cause clogging of the size exclusion column.

**53.** Save a 50 µL aliquot of the filtered dialyzed eluate.

**54.** Manually inject 5 mL of the filtered dialyzed eluate onto the HiLoad 16/600 Superdex 200 pg column.

**55.** Carry out the size exclusion chromatography (SEC) with 1.5 CV of 1x SECB buffer at a flow rate of 1 mL/min, collecting 0.8 mL fractions from 40 mL to 110 mL elution volumes in a 96 deep-well plate.

Pause point: The SEC fraction plates can be safely stored overnight covered with Parafilm M at 4 °C.

**56.** The A280 profile of the run should show three distinct peaks at approximately 48.1 mL, 71.6 mL and 97.9 mL volumes. Peak 2 at ≈ 71.6 mL corresponds to the desired 10His-pG-Tn5 homodimer eluting at ≈ 127 kDa.

**57.** Analyze every fraction of the peak 2 by SDS-PAGE with Coomassie staining, to determine which fractions contain significant amounts of cleaved 10His-pG-Tn5:Tn5 heterodimer. Load ≈ 10.5 µL per lane of 0.8 mL SEC fractions corresponding to peak 2. The inflexion of the peak 2 shoulder at elution volumes above 71.6 mL will contain cleaved 10His-pG-Tn5:Tn5 heterodimer which is smaller than 127 kDa.

**58.** Based on the results of the SDS-PAGE, decide which SEC fractions to pool. We typically pool 11 0.8 mL SEC fractions to give 8.8 mL SEC pool.

Note: It is best practice to be stringent about excluding SEC fractions that visibly contain cleaved 10His-pG-Tn5, and to prioritize stock purity over total yield. However, it is normal for a small degree of 10His-pG-Tn5 cleavage to be seen on SDS-PAGE which is induced during sample denaturation, this can be limited by denaturing samples in loading buffer at 70 °C for 10 minutes.

**59.** Save a 100 µL aliquot of the SEC pool.

**60.** Measure the protein concentration of the SEC pool using a Bradford reagent kit, a fluorescence-based Qubit protein kit or a Nanodrop spectrophotometer A205 measurement.

Note: Different protein concentration determination methods are likely to give discrepant measurements due to the composition of the 1x SECB buffer. We do not typically use Nanodrop A280 readings as a reliable method for measuring protein concentration. Instead, we dilute an aliquot of the sample 5-fold in 50 mM Tris-HCl pH 7.5 and measure 2.0 µL of this diluted sample using the Qubit protein kit, then back-calculate the protein concentration in the original sample.

**61.** Determine the volume to which the SEC pool sample must be concentrated to in order to give a final concentration of 2 mg/mL. Multiply the determined protein concentration of the SEC pool by its total volume and divide by 2 mg/mL to give the target volume to concentrate down to. Example: 8.8 mL SEC pool at 0.50 mg/mL = 4.4 mg total protein, divided by 2 = concentrate down to 2.2 mL to achieve a final concentration of 2 mg/mL.

**62.** Pre-equilibrate a 15 mL sample-capacity 30 MWCO Amicon Ultra concentrator with 5 mL 1x SECB, spinning at 4’000 xg at 4 °C. Note down the time it takes for the buffer to completely pass through the filter, which should take about 4 minutes.

**63.** Remove and discard the 1x SECB buffer flowthrough in the lower chamber of the Amicon Ultra concentrator.

**64.** Transfer the entirety of the SEC pool sample to the pre-equilibrated Amicon Ultra concentrator, and spin for 2 minutes at 4’000 xg at 4 °C.

**65.** After the 2 minutes spin, check the volume of the sample flowthrough, to get an idea of the length of time it will take to concentrate the SEC pool to your desired volume.

**66.** Using a P1000 pipette, gently pipet the SEC pool sample in the upper Amicon Ultra chamber, being careful not to stab the membrane or introduce bubbles which can denature the 10His-pG-Tn5 protein. This step helps to prevent the protein from ‘crashing out’ and becoming insoluble, which is caused by a local increase of protein concentration close to the filter membrane.

**67.** Continue to spin in 2-minute increments or less, repeating the P1000 pipetting with each spin.

**68.** Once the SEC pool sample has been concentrated to the desired volume, measure the protein concentration again, verifying the sample is at ≈ 2.0 mg/mL.

**69.** Heat an aliquot of 100% UltraPure glycerol to 50 °C to aid in accurate pipetting.

**70.** Pipet out a volume of 100% UltraPure glycerol equal to your 2.0 mg/mL 10His-pG-Tn5 stock, to a fresh empty tube of appropriate volume, and place on ice to chill prior to adding 10His-pG-Tn5.

Warning: Accurate pipetting is critical for long term stability of the stock, as it affects the final storage buffer composition. Make sure to accurately measure both the exact volume of the SEC pool concentrate as well as the 100% UltraPure glycerol. Make sure to sufficiently chill the aliquot of heated glycerol on ice prior to proceeding to combine with protein.

**71.** Once the glycerol has cooled on ice, add the SEC pool concentrate containing 10His-pG-Tn5 to the glycerol, note the difference in refractive index between the glycerol and the protein. Mix by gentle pipetting with P1000, avoiding creating bubbles. Mix very thoroughly until glycerol is immiscible.

**72.** Measure the protein concentration of the 10His-pG-Tn5 stock, which should be 1 mg/mL.

Note: The final concentration of 55% glycerol in the stock solution can interfere with accurate protein concentration measurements, therefore perform any measurements with several dilutions of the stock and back-calculate the concentration as necessary.

**73.** Make 1 mL aliquots of the glycerol stock of 10His-pG-Tn5 in Eppendorf 1.5 mL Protein LoBind tubes.

**74.** Store the 10His-pG-Tn5 stock aliquots in an enzyme cooler block at –20 °C, to minimize temperature fluctuations. When stored under these conditions, we have found 10His-pG-Tn5 is stable for > 21 months without noticeable loss of activity.

Warning: Do not store the glycerol stocks at –80 °C. Do not load Tn5 with MEDS adapters prior to storage at –20 °C, as doing so changes the final storage buffer composition and causes loss of Tn5 activity. Only load small, single-use aliquots of Tn5 with adapters, as detailed in Section B of this protocol.

---

#### OpenTn5 Protocol

##### Section B: λ gDNA Tagmentation Assay

Here we describe a rapid and sensitive method for quantification of DNA fragmentation by transposition, referred to as the tagmentation activity of Tn5. For regular applications such as ATAC-seq and CUT&Tag Nextera Mosaic End Double Stranded (MEDS) adapters can be ordered as desalted duplexes from IDT or ordered as ssDNA oligos and annealed in the lab. For more sensitive applications, such as single cell assays, we recommend ordering PAGE-purified duplexes. Note that the 5’-phosphorylation on the bottom MEDS oligo is essential for tagmentation activity.

The predicted molecular weight of a pG-Tn5 monomer is 63.4 kDa. Therefore, a 1 mg/mL stock has a 15.8 μM concentration. A single monomer of pG-Tn5 binds one MEDS adapter molecule. The $K_d$ of $Tn5^{E54K, L372P}$ for MEDS DNA has been measured to be ~ 0.50 μM by Hennig et al. Under the conditions of our assay, the sub-stoichiometric amounts of MEDS adapters are expected to be fully bound by Tn5. We use lambda phage genomic DNA (λ gDNA) purchased from NEB as the substrate DNA for tagmentation. Under the conditions of our assay, the λ gDNA is limiting and all the available assembled MEDS-pG-Tn5 transposomes are used up in the reaction. Tagmentation is a single-turnover reaction, resulting in the MEDS adapters covalently transposed into the template DNA. Post-transposition the Tn5 dimer remains tightly bound to the template DNA and must be subsequently stripped off using SDS for downstream analyses.

##### Nextera MEDS adapter oligo sequences

| | Sequence 5'–3' | |
|---|---|---|
| Nextera N5 adapter strand 1 | TCG TCG GCA GCG TCA GAT GTG TAT AAG AGA CAG | Order as a duplex |
| Nextera N5 adapter strand 2 | /5Phos/CTG TCT CTT ATA CAC ATC T | |
| Nextera N7 adapter strand 1 | GTC TCG TGG GCT CGG AGA TGT GTA TAA GAG ACA G | Order as a duplex |
| Nextera N7 adapter strand 2 | /5Phos/CTG TCT CTT ATA CAC ATC T | |

1. Make a working stock by diluting an aliquot of 100 μM stocks of each P5 and P7 Nextera MEDS adapters to 20 μM, in 10 mM Tris-HCl pH 7.5, 0.1 mM EDTA (IDTE).

2. Make a working stock of MEDS adapters by combining equal volumes (1:1) of 20 μM stocks of P5 and P7 adapters. This adapter stock has a 10 μM concentration in terms of each P5 and P7 adapter but remains 20 μM in respect to the MEDS DNA. Since it is the MEDS sequence that is bound to by Tn5, we refer to this working stock as 20 μM adapters.

3. Make a dilution series of 20 μM adapters in IDTE, 50 μL each, 16, 14, 12, 10, 8 and 6 μM.

4. Combine 10 μL of pG-Tn5 stock (55% glycerol) with 10 μL of adapters for each of the dilutions and mix by gentle pipetting.

5. Assemble the transposomes by incubating at 25 °C for 10 minutes in a thermocycler. Transposomes can be kept at room temperature for the duration of assembling the activity assay.

Note: Transposomes can be stored short term at 4 °C or –20 °C for 1 to 2 days but lose activity quickly under these conditions and are not suitable for long term storage. Therefore, assemble transposomes fresh as required on the day of experiment. We have found that adapter loading at RT on the bench for 30-60 minutes gives similar results to incubating at 25 °C for 10 minutes in a thermocycler.

6. Prepare a mastermix of λ gDNA in TD buffer, account for a no transposomes control and include a 2-reaction excess.
Note: λ gDNA is viscous due to its large molecular weight (48.5 kb), excessive vortexing and freeze-thaw cycles may induce fragmentation over time. To alleviate this, make small 20 µL aliquots of the NEB stock to store at –20 °C. Once thawed, store λ gDNA at 4 °C

7. Aliquot 21 µL of the reaction mastermix into individual PCR strip tubes, one for each transposomes loading condition and one for no transposome control. Carefully pipette 4 µL of each transposome of the MEDS dilution series into the pre-aliquoted λ gDNA in TD buffer. Pipette gently to mix.

8. Incubate the tagmentation reaction at 55 °C for 10 minutes in a thermocycler.

9. Quench the reaction by adding 1.5 µL of 4% SDS, vortexing well until each sample is foamy.

   Note: Failure to inadequately quench the reaction will result in Tn5 remaining to be bound to λ gDNA post-tagmentation and give rise to anomalous migration of DNA in agarose gels and Tapestation Screentape. Inadequate quenching can be diagnosed by a lack of foaming after SDS addition, as well as DNA remaining to stay in the agarose gel well.

10. Quick-spin the samples in a table-top microcentrifuge

11. Analyze 1 µL of the quenched reactions, equivalent to 9.4 ng λ gDNA, on a genomic DNA Screentape in a Tapestation instrument.

    Note: The measured DNA concentration of tagmented DNA will be higher than this due to the incorporation of the mass of MEDS adapters into the tagmented template DNA.

    The size of the tagmented DNA should decrease as a function of increasing [MEDS]. Sufficiently tagmented λ gDNA should produce a major band at ~250 bp. Complete tagmentation of λ gDNA should occur readily at close to equimolar concentration of MEDS, if all pG-Tn5 protein is native in functional homodimers. At 1 mg/mL the pG-Tn5 stock has a 15.8 µM concentration, each Tn5 molecule binds one MEDS adapter. Assuming all protein is present in functional dimers, pG-Tn5 should be saturated at 16 and 20 µM adapter loading ratios. However, as the exact fraction of correctly-annealed adapters in the stock solution can vary, the exact loading ratio needs to be determined empirically.

    When the available Tn5 has been saturated by MEDS adapters, free MEDS adapters will be present as a ~50 bp band. We typically include a slight molar excess of MEDS based on the activity assay, as under-loading Tn5 results in a non-linear size increase shift of tagmented

DNA product. However, addition of a large excess of MEDS results in the free adapters compete with substrate DNA for Tn5 binding and generates poor quality sequencing libraries.

**Comparison of Tagmentation Activity to Illumina TDE1 Enzyme**

We define the tagmentation activity of Tn5 in terms of a Tagment Units (TU). One Tagment Unit of Tn5 is defined as the amount of MEDS adapter-saturated Tn5 required to uniformly tagment 125 ng of λ gDNA to < 250 bp size (effective size including the incorporated adapters). This corresponds to 1 µg of 10His-pG-Tn5 homodimer saturated with MEDS adapters, or 2 µL of fully assembled pG-Tn5 transposome at 0.5 mg/mL. We have determined the concentration of Tn5 in Illumina TDE1 enzyme to be around 1/16 of the pG-Tn5 stock, or 0.0625 mg/mL, based on SDS-PAGE Coomassie staining. Illumina TDE1 enzyme shows correspondingly 8-fold less tagmentation activity on λ gDNA when compared to an equal volume of assembled pG-Tn5 transposome at 0.5 mg/mL. Therefore, 8-fold less volume of pG-Tn5 transposome at 0.5 mg/mL may be used to replace Illumina TDE1 enzyme, for example 0.31 µL pG-Tn5 replacing 2.50 µL TDE1 in a standard 50’000-cell OmniATAC reaction.
